# 3D dynamic multiscale force and shape analysis of *in-vivo* elastic stress sensors

**DOI:** 10.1101/2025.01.22.633835

**Authors:** Alejandro Jurado, Jonas Isensee, Arne Hofemeier, Lea Johanna Krüger, Raphael Wittkowski, Ramin Golestanian, Philip Bittihn, Timo Betz

**Affiliations:** Third Institute of Physics-Biophysics, University of Göttingen, Göttingen, Germany; Cluster of Excellence “Multiscale Bioimaging: from Molecular Machines to Networks of Excitable Cells” (MBExC), University of Göttingen, Göttingen, Germany; Institute of Pharmacology and Toxicology, University Medical Center Göttingen, Göttingen, Germany; Max-Planck-Institute for Dynamics and Self-Organization, Göttingen, Germany; Institute of Theoretical Physics, Center for Soft Nanoscience, University of Münster,Münster, Germany

**Keywords:** Sotfware, Biomechanics, Force Inference, In-vivo, Multi-scale

## Abstract

The measurement of stresses and forces at the tissue level has proven to be an indispensable tool for the understanding of complex biological phenomena such as cancer invasion, embryo development or wound healing. One of the most versatile tools for force inference at the cell and tissue level are elastic force sensors, whose biocompatibility and tunable material properties make them suitable for many different experimental scenarios. The evaluation of those forces, however, is still a bottleneck due to the numerical methods seen in literature until now, which are usually slow and render low experimental yield. Here we present BeadBuddy, a ready-to-use platform for the evaluation of deformation and stresses from fluorescently labelled sensors within seconds. The strengths of BeadBuddy lie in the pre-computed analytical solutions of the elastic problem, the abstraction of data into Spherical Harmonics, and a simple user interface that creates a smooth workflow for force inference.

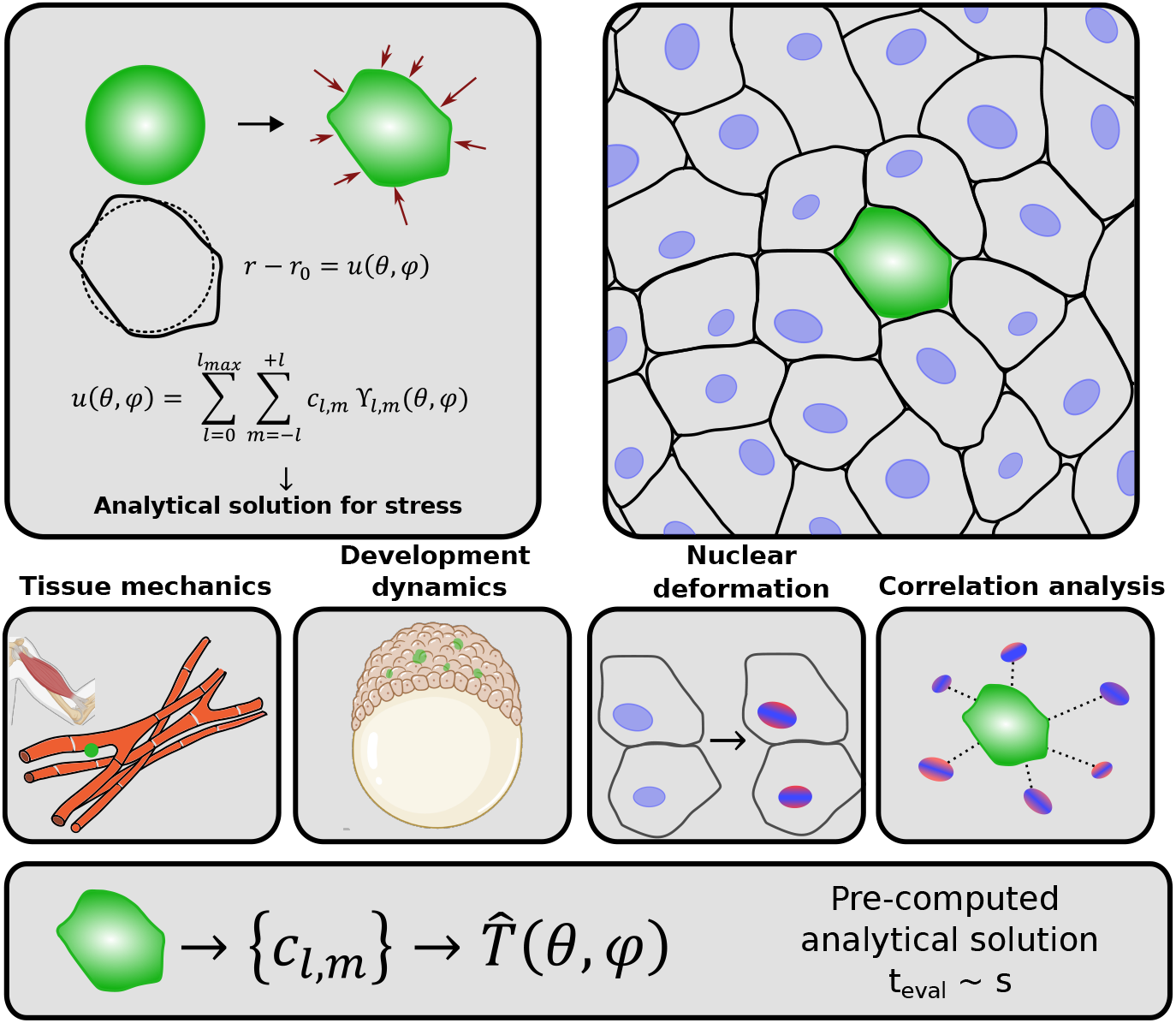

## 1 Introduction

The field of **cell and tissue mechanics** is rapidly evolving, with new methods and measurement techniques being proposed at a fast pace, and new goals of ever increasing difficulty and interest appearing on the research horizon [1–4]. Although cell and tissue biochemistry is instrumental for the understanding of many key biological phenomena such as wound healing, embryogenesis or cancer invasion, a full, comprehensive picture additionally needs to integrate the cell mechanical aspects, which complement and reinforce the models. The consideration of physical parameters such as adhesion forces [5], surface tension [6], viscosity [7] or elastic energies [8] has proven to be a necessary and beneficial complement to the existing models, casting light onto many yet unexplored biological mechanisms. This idea is at the very core of the field of mechanobiology, creating an integrated view in which gene expression, chemical pathways and protein folding come together in models with adhesion forces, elastic moduli or strain measurements.

Of particular interest is the possibility of measuring forces at the cell scale, which is provided by a series of *in vitro* approaches, such as Pipette Aspiration, Atomic and Traction Force Microscopy, or Optical Tweezing (see reviews in [9, 10]). In contrast, while *in vivo* experiments are much more powerful due to their holistic nature, similar force interference techniques are not developed to their fullest potential, as these often remain destructive, render a low statistical yield, or are limited to surface investigation such as Laser Ablation [11] or Microindentation [12]. The potential to overcome these weaknesses even *in vivo* is provided by micro-beads as force sensors, which are injected directly into living tissue [13, 14] and organoids [15, 16]. This is a relatively new technique which has gained much popularity in the last years.

Since the pioneering work of Campàs et al. in 2014 using oil droplets [17], many groups have replicated this experimental technique, improving the injection methods, the fabrication of droplets [18] and experimenting with new, more versatile and tunable materials, like PDMS [19] and ferro-fluid materials [20]. The general advantage of this approach relies on the fact that micro-droplets are innocuous for the organisms, and the experiments can be done in such a gentle manner that systems as fragile as a developing embryo can be studied without introducing artifacts. Given that the mechanical properties of the injected material are known, the deformation inside the tissue can be related to the force felt by the sensor. However, this step is cumbersome and difficult to accomplish if one lacks the proper mathematical and computational tools, and still remains the bottleneck for most labs who want to try out this technique. The analysis of deformed beads or droplets as force sensors has been performed previously using different approaches. In the initial work of Campas et al. [17], Laplace’s law relating the local curvature of the oil beads with local surface stresses was used. In a different approach of the same group, the ratio of the main axes of a fitted ellipsoid were analyzed [20], simplifying the problem to a one-dimensional deformation. Similarly, gel sensors have been used to measure local stress changes in tissue based on isotropic volume changes [21]. In more complex approaches, elastic hydrogel bead deformation has been analyzed using finite element (FE) approaches [14] and more recently, numerical optimization based on analytical solutions has been applied [22]. Even though these works have immensely contributed to the field, paving the way and showing the potential of the technique, a straightforward analysis system – ideally with a simple user interface - is still missing for a full adoption of the technique.

To overcome this hurdle we present **BeadBuddy**, a ready-to-use platform that directly takes the microscope images as input and provides a graphical user interface to not only obtain the deformation and surface stresses from the force sensors, but also to intrinsically separate these into the relevant length scales at which they are acting. To construct the working frame of BeadBuddy, we combined the powerful segmentation library PyClEsperanto [23] with a decomposition into Spherical Harmonics (SH) of the data [24] and pre-computed analytical solutions for the elastic problem [25]. In its compact form, BeadBuddy allows **fast batch analysis of force sensors** inside tissue, pushing forward the rapid acquisition of statistics *in vitro* or *in vivo* tissue. The framework can also be applied to segment and deliver a 3D shape analysis in terms of spherical harmonics for other objects, such as nuclei or liquid droplets.

The necessary steps to achieve this goal are presented using a developing zebrafish embryo as showcase example. The first sections of this article are aimed to guide the reader through the analysis steps of the process which constitutes the core of BeadBuddy. The last section is an endeavour to show several particular analysis examples which any user could tackle easily and in a matter of minutes thanks to our software.

## 2 Data processing

Starting from the acquisition of fluorescent images in the microscope, there are several steps that an analysis pipeline must cover in order to properly analyze the deformation of small objects at the cell scale. These steps include a proper image segmentation, an abstraction of the measured geometries into a compact representation, the mathematical solution of the elastic problem, and the post-processing of the obtained data to achieve physical magnitudes from which one can construct useful statistics. The following subsections will guide the reader through the steps that lead us to construct BeadBuddy as a stand-alone software.

### 2.1 Image segmentation

Segmentation is the first step in almost any analysis of fluorescently labelled images, which in the context of shape analysis has to focus on separating the signal from the background as cleanly and accurate as possible. We constructed our segmentation pipeline based on the versatile library PyClEsperanto [23], and optimized it to detect small fluorescent objects over a dark background. To showcase the flexibility of BeadBuddy beyond its application in tissue stress inference, we first highlight its capabilities in image segmentation and shape detection using nuclei of developing Zebrafish as an example. We segmented the fluorescently marked nuclei of an embryo at 7 hpf (hours post fertilization), which served as a benchmark and quality control of our pipeline. Since an embryo at this early stage contains thousands of nuclei where many are hidden under several tissue layers, nuclei segmentation and shape analysis is a difficult task. Thanks to the high penetration of our Light Sheet Microscopy technique, the isotropic resolution achieved after a MultiView fusion and the flexibility of our segmentation pipeline, the nuclei in the embryo can be segmented with high precision, as can be seen in Figure 1A. Comparing with a manual count of the same dataset, BeadBuddy could detect 94% of the nuclei present in the embryo at 7 hpf.

**Fig. 1.**
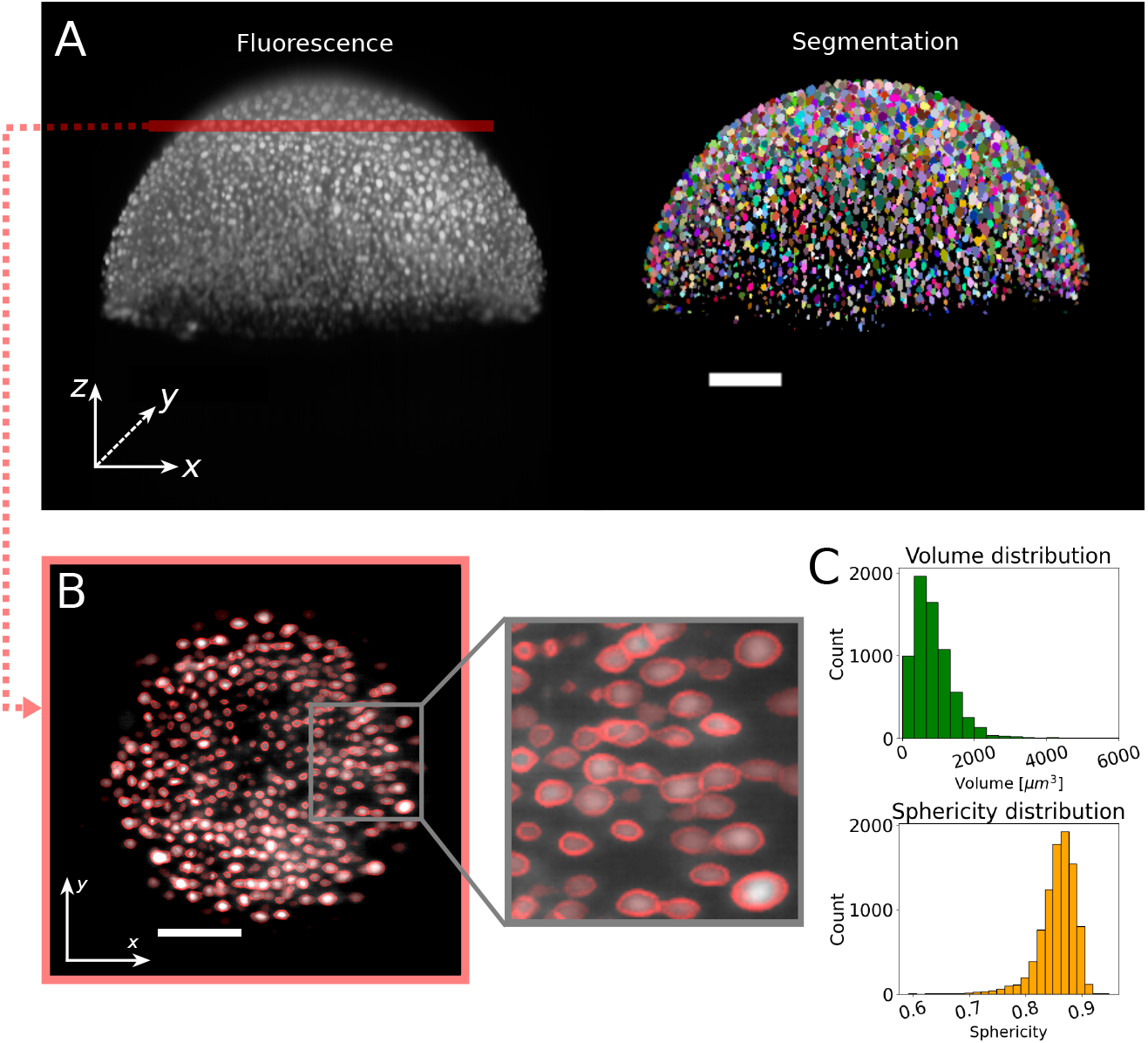
Output of an optimal segmentation of lightsheet data. **(A)** 3D acquisition of H2AmCherry nuclei in a 7 hpf Zebrafish (left) and its segmentation using BeadBuddy (right). **(B)** Cross section of the embryo through the red line in (A). The outline of each segmented object (red) is compared to the fluorescent data (grayscale) to assess the quality of the segmentation. **(C)** Distribution of volumes and sphericities of the segmented nuclei. Scalebars 100 µm.

The quality of the segmentation can be visually verified (Figure 1B) by checking whether the majority of the fluorescent bodies have been detected, and can be dynamically corrected in cases of under- or oversegmentation. BeadBuddy offers several **segmentation parameters** for the user which control the expected size of the bodies, their relative brightness and their closeness (see tutorial in Appendix A). An initial set of parameters is suggested after software initialization which have been proven to work in several microscopy and tissue combinations. These parameters can be then used as a starting point for fine-tuning alone.

Already from this segmentation some relevant information from the objects can be extracted with minimal post-processing. For some biological problems, it is usually sufficient to obtain general geometrical information, such as the volume of the found objects, their spatial distribution (centers of mass, density) or their sphericity. In Figure 1C, histograms of two of these parameters can be seen, namely volume and sphericity, defined here as the ratio of the surface area of a sphere with the same volume to the surface area of the object,

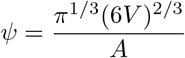

with *V* and *A* the volume and surface area of the object.

In our particular example, we find a typical volume centered around 450 µm^3^, corresponding to nuclei with diameter of ≈ 9.5 µm. From the segmentation, also the sphericity of an object can be computed, which can help to illustrate the extent of deformation of a spherical object. In this particular case, Zebrafish nuclei at 7 hpf exhibit a distribution of sphericity centered around *ψ* = 0.86. Even though these measurements can already be quite useful for analysis and to better understand tissue morphology and dynamics [19, 26, 27], it barely scratches the surface of possibilities. In order to take the analysis a step further, we need to transform the segmentation data into a different representation space. We aim to use a mathematical base which contains information related to the different length scales of the deformations, and from which stress can be inferred. Following the example of previous similar analyses [28–30], we opt for an expansion of each detected body into SHs. This representation is useful here because it provide an intrinsic length scale separation by their degree, and furthermore, they present an orthonormal set of basis functions solving the Laplace equation.

### 2.2 Spherical Harmonics representation

SHs are a set of functions ϒ *(θ, ϕ)* which emerge from the solution to the angular part of Laplace’s equation, one of the most fundamental physical relationships, appearing in the study of electrostatics, quantum mechanics or fluid mechanics, to name some. The great advantage of SHs is that, due do this general nature, they can be used in an expansion to express any well-behaved function *u(θ, ϕ)* in the spherical domain *θ* ∈ [−π/2, π/2], *ϕ* ∈ [−π/2, π/2] such as a body deformation

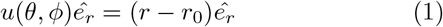

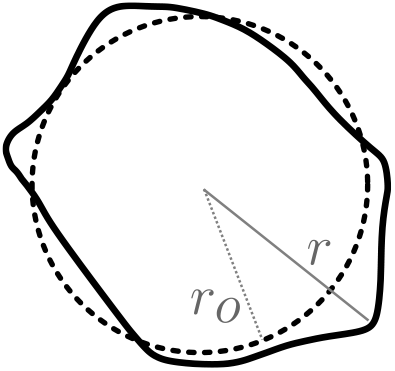

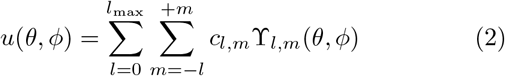

where *θ* and *ϕ* are polar and azimuth angles, *ê*_*r*_ is the unitary radial vector in spherical coordinates, ϒ_*l,m*_ the SH functions and c_*l,m*_ the coefficients of the expansion for each pair of degree and order (*l*, *m*). Please note that the naming convention for”degree” and “order” may be different in other fields. Here we stick to the most common mathematical convention, where *degree* preceeds *order*. The definition of Spherical Harmonics used in this work can be found in Appendix B.

The intuition to expand the surface of any concave and simply connected object in 3D onto this base emerges naturally. Since the functions ϒ (*θ, ϕ*) are known and well defined, this representation brings the great benefit that the information of a complex volumetric structure is encoded in a table of coefficients {*c*_*l,m*_}.

On the one hand the coefficient encoding results in a **drastic reduction of data** that has to be handled. As will be seen later, the expansion of a micrometer-sized body such as a zebrafish nucleus maintaining micron resolution can be achieved with degree *l*_*max*_ *≈*9, amounting to no more than a hundred coefficients. On the other hand, **the hierarchical separation into length scales is intrinsic** to SHs, with each degree *l* carrying information equivalent to that of a wavelength, or typical deformation size. This is particularly useful when analyzing dynamical data as single degrees can be followed separately:

- **Low Degree Deformations** are equivalent to changes in volume (*l* = 0), translations (*l* = 1) and prolate/oblate approximations (*l* = 2), carrying information about global features, such as compression/expansion and main elongation directions. Particularly useful are the parameters |*c*_2,*m*_ |/*c*_0,0_, used previously in some studies [13, 15] to indicate the main deformation dipoles.
- **High Degree Deformations** contain information about local features, such as deformations due to cell-cell or organelle-nucleus interactions. If the microscopy technique being used offers enough resolution, or the studied objects are big enough, high-order components can contain information about important phenomena on a finer scale, such as membrane undulations or wrinkling as well as geometries related to strong confinement (sharp edges, straight interfaces).

Starting from the detected surface of the segmentation, an optimal expansion is found using the python-based library ***SHTools*** [24]. Figure 2 shows the power spectrum of the decomposition alongside the shapes resulting from a truncation at different degrees *l*_max_. It can be observed how the expansion to degree *l*_*max*_ = 2 is simply an ellipsoid roughly oriented in the direction of maximum elongation of the body. The higher the maximum SH degree, the more refined the surface becomes, attaining smaller features of deformation. The power spectral density of each order, computed as

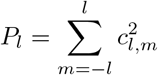

helps to identify the dominating modes of deformation and their distribution. In Figure 2B a simplified illustration of the modes ϒ_*l,m*=*l*_ is shown below the SHs spectra, exhibiting the corresponding deformation wavelength each mode carries.

**Fig. 2.**
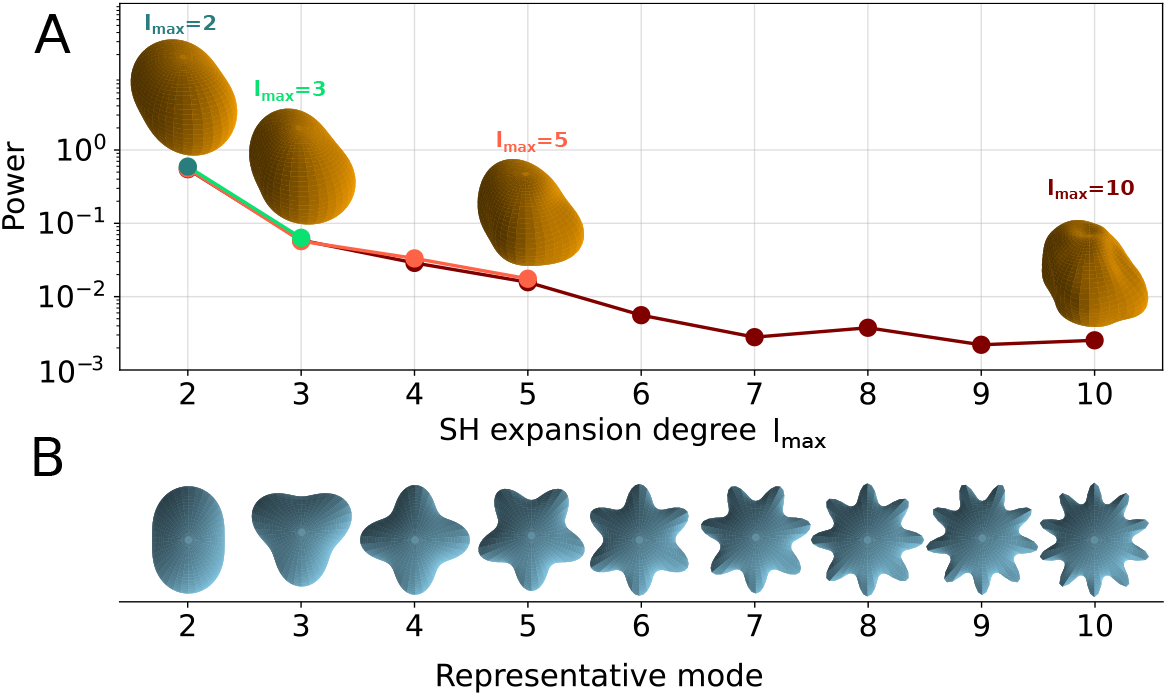
Expansion of a measured body using different SHs truncations. **(A)** Spectra of the expansion of an irregular body into SHs, with increasing maximum degree *l*_max_. Each degree captures more details of the surface. Different matching colors show the spectrum of each expansion. **(B)** Representative modes of Spherical Harmonics ϒ_*l,m*=*l*_ displaying their typical undulation lengthscale.

To gain an understanding of the corresponding lateral resolution related to each mode, we define the node-antinode distance *d*_*NA*_. With *r*_0_ the radius of the initial, undeformed spherical bead, and imposing volume conservation in the bead, it holds that

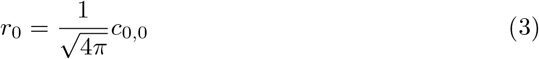

from which the distance between a node and an anti-node of the *l*^*th*^ mode is

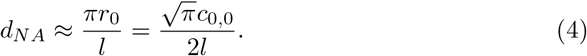

This approximation tells us that for a typical bead or nucleus of radius 5µm, a maximum degree of expansion *l*_max_ = 10 would render a lateral resolution of ≈ 1.6*µm*. This must be, nevertheless, compensated by the available resolution and the amount of points in the surface detection. Whenever the amount of SH coefficients for a particular fit exceeds the amount of available surface points, we are effectively overfitting the available data. This is reminiscent to the Nyquist frequency in Fourier decomposition. We included a sanity test in BeadBuddy to only allow fits which satisfy this condition, as to not introduce artifacts in the analysis.

### 2.3 Analytical solution to the elastic problem

The mathematical representation in SHs is well suited for dealing with mathematical problems in spherical geometries, since they are a set of orthogonal functions and can be used as a basis for constructing more complex mathematical structures. In the particular case of force sensors in living tissue, one can apply linear elasticity as long as the elastic deformations are considerably smaller than the scale of the objects, which we necessarily consider to hold true to tackle our problem. Using Hooke’s law in three dimensions [31], the relationship between stress 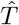 and strain 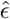 is

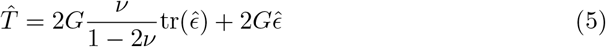

where the coupling terms are the Poisson ratio *ν* and the shear modulus *G*, and 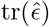 stands for the trace of the strain. The relation between strain and deformation 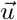 is

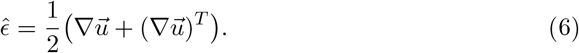

Assuming equilibrium, the divergence of the stress tensor 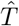 is equal to the external forces per unit volume [25, 31]. Since we assume that the external forces are negligible compared to the stress exerted on the bead surface, it follows that 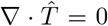 in the bulk and one comes to the fundamental relationship

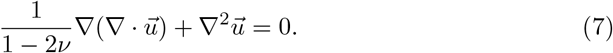

To retrieve the stress responsible for a given change of shape of a sphere, the deformation needs to be expressed properly for every point of the surface. However, given the nature of typical fluorescence data, the deformation field is not uniquely determined by the surface, since many different deformations could lead to the same shape. This makes a choice necessary: one can formulate the problem with a completely unrestricted deformation, which leads to an a computationally expensive optimization problem (see [22]), or we can work around it assuming that tangential deformations are negligible, and only considering radial deformations. It should be mentioned that inert stress sensors, on which cells cannot apply any tangential forces, nicely fulfil this constraint

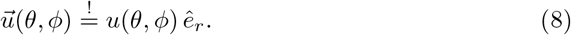

This latter approach allowed us [25] to fully solve the stress tensor analytically with a Spherical Harmonics ansatz. Using this approach, the radial displacement of an object can be expanded in SHs up to an arbitrary degree, and the stress tensor is then fully determined by the expansion coefficients and the mechanical properties of the material:

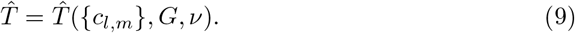

The default version of BeadBuddy includes pre-computed analytical expressions to quickly evaluate the solutions to the elastic problem up to a degree *l*_*max*_ = 15, with the possibility of solving and storing higher-order solutions for future use. The great advantage of this method relies on the pure analytical nature of the calculations, for which only the coefficients of the SH expansion are needed. Once the coefficients are known, they are substituted in the stored template expressions, effectively rendering solutions on the order of seconds for a typical office computer. In order to achieve this speed, we sacrifice the consideration of tangential displacements of the surface of the sensors. While this seems to be an important downside, many elastic hydrogel particles are chemically inert, meaning that cells cannot apply any pulling or tangential forces. For such situations the here proposed method is optimized and sufficient.

To test the accuracy of our method against other well established methods, we compared BeadBuddy solutions against COMSOL-based finite element stress inferences using the same deformation fields as in the analytical approach. A direct comparison of the radial stress field is shown below.

Figure 3A shows a direct comparison between our analytical solution and a standard finite element solution. The discrepancies did not show a systematic pattern and this feature was reproducible for the 30+ beads that were used in the comparison, which leads to the question of whether the differences arise from the truncation of the analytical solution to a maximum degree *l*_max_. If this were the case, it would be reflected in the spectral decomposition of the difference field in Figure 3C. As can be observed, higher degrees do not carry a particularly high contribution, implying that the discrepancy does not emerge exclusively from the truncation, but might also be related to the assumption of only radial displacements. In order to directly compare the forces *F*_BB_ corresponding to BeadBuddy and *F*_COMSOL_ corresponding to the FE calculations, the ratio *F*_COMSOL_/*F*_BB_ was calculated for several degrees *l*_max_ of the analytical solution, as seen in Figure 3B. Interestingly enough, the best agreement was achieved for analytical solutions of degree *l*_*max*_ = 5, further reinforcing the idea that solution discrepancies do not necessarily improve increasing the maximum expansion degree *l*_max_ analytically. The inset in Figure 3C shows the computation time for different order of the analytical solution, which for degree *l*_max_ = 6 is barely above one second. This is a major improvement as compared to FE computation for the same beads as implemented in COMSOL, requiring ∼ 20 s per bead in our tests.

**Fig. 3.**
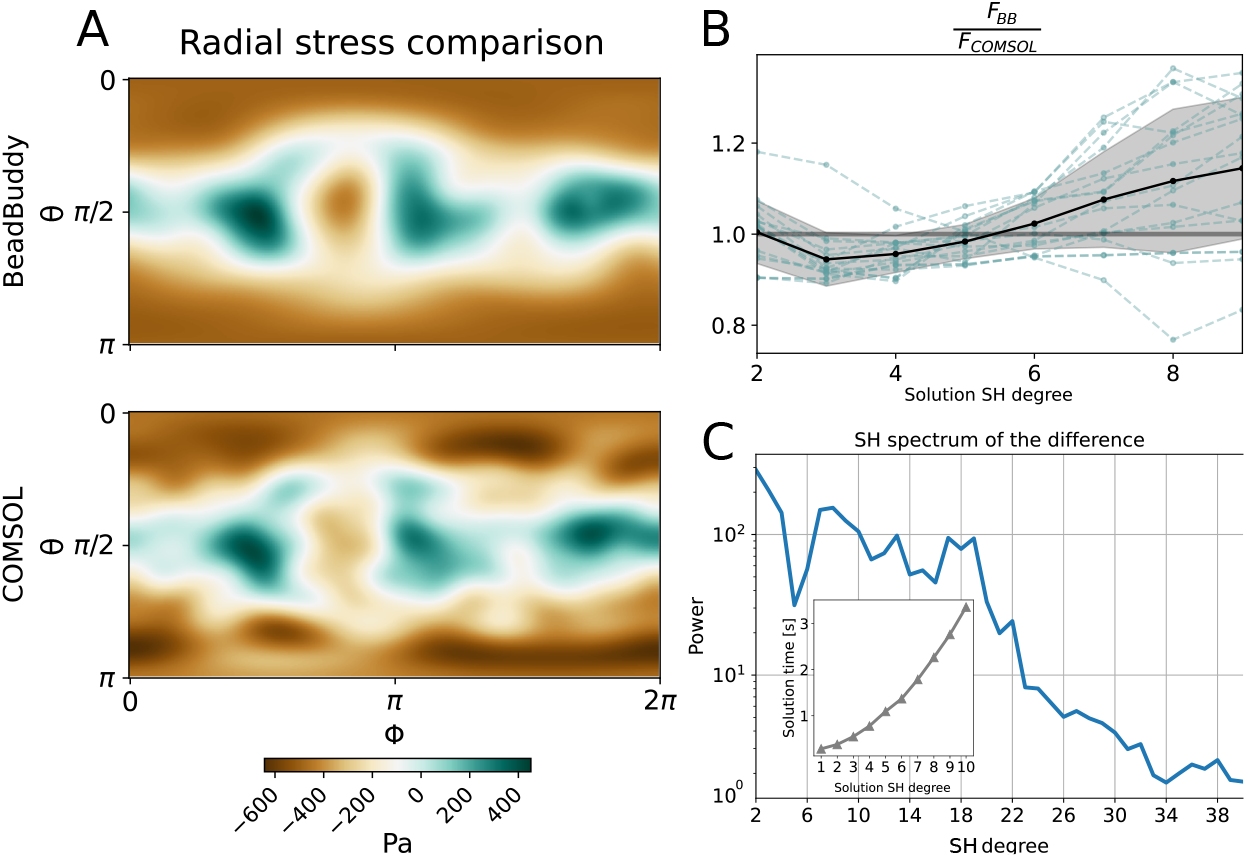
Benchmark of BeadBuddy’s analytical solutions accuracy against a finite element approach. **(A)**. Plane projections of the stress map as calculated on the surface of a body by BeadBuddy with expansion degree *l*_max=5_ (upper) and COMSOL (lower). **(B)** Ratio of the numerically integrated force over the surface for the analytical solution and FE solution. The dotted lines show the solutions for individual beads and the black line shows the average ratio, along with its standard error in grey. **(C)** Power spectral density of the absolute stress difference between BeadBuddy and COMSOL, calculated for the stress map in A. The inset shows the average solution time of BeadBuddy for a truncation degree *l*_max_.

## 3 BeadBuddy in action: Detailed examples and case studies

Having the basic working principles of BeadBuddy established, we showcase its unique features by applying it to a series of common examples. Out of the data extracted with BeadBuddy in its **batch analysis mode**, many different studies can begin based on the distributions of volumes, deformations, SH spectra with the implied length scales, and the associated stresses and forces. In the following we demonstrate the flexibility of the data outputted by BeadBuddy. We start with nuclei and force-sensor analysis in Zebrafish, before moving to a reconstituted muscle tissue, in which the distribution of forces reveals itself as a consequence of the tissue architecture.

### 3.1 Nuclear deformation in developing Zebrafish

The correlation between nuclear deformation and the stress acting on the cells is widely discussed in the literature. Several studies have hinted to the fact that the cell nucleus might be a central coordinator of mechanotransduction, and how its shape and mechanical properties are directly linked to the cell activity and function [32–35]. Even though the mechanical properties of nuclei are rapidly changing in a tissue and can not be reliably measured on such scale, the analysis of the deformation characteristics might already shed some light on mechanical patterns in tissue.

The batch analysis capabilities of BeadBuddy allow the study of several thousands nuclei in a matter of minutes. The reduction of data from cumbersome 3D fluorescence images to a well structured relation of geometrical parameters thanks to the expansion in SHs allows for such an **a-priori study of mechanical patterns**. We ask the question of whether nuclear deformation patterns can be found in the early stages of a developing Zebrafish embryo. To simplify the analysis, BeadBuddy can find the optimal rotation for each shape which minimizes the SH coefficient c_*l*=2,*m*=0_, orienting the body in such a way that shows its shortest axis along the z axis. In this orientation, a convenient deformation parameter is defined as:

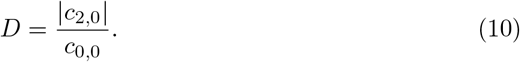

Note that the decision to use *c*_*l*=2,*m*=0_ is arbitrary, and any harmonic component from the second degree would have produced the major deformation direction after its minimization. In this respect, this analysis is equivalent to the oblate/prolate fit of a body. As shown in Figure 4A, the value D = 0.2 is a good threshold for marking strong deformations. This helps with the classification of nuclei in this particular tissue. A schematic of spherical deformation is included in Figure 4A to create an intuition of the geometry associated to this threshold.

**Fig. 4.**
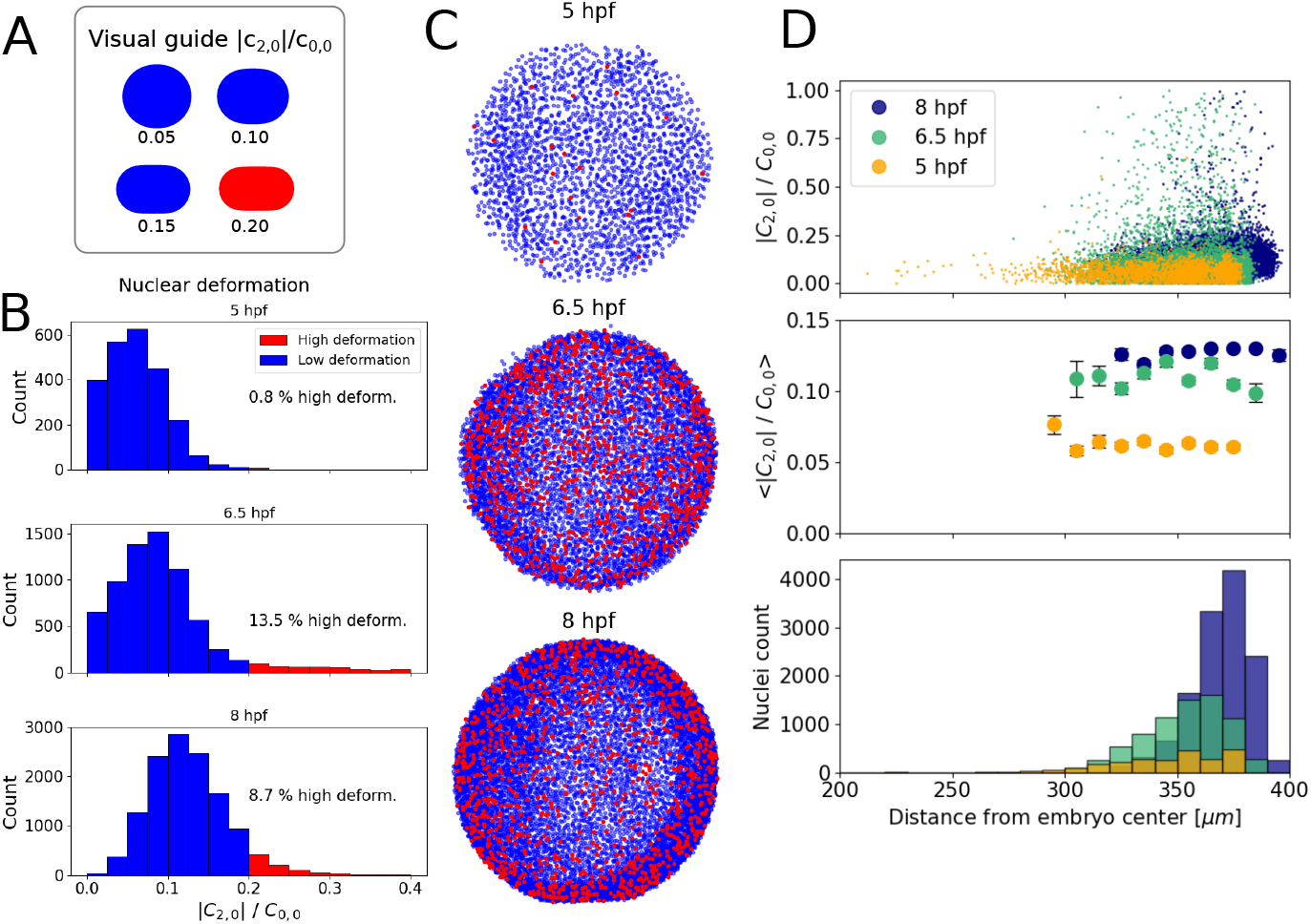
Study of nuclear deformation distribution in a developing Zebrafish embryo. **(A)** Schematic depicting the side view of a sphere which has been deformed with increasing value of *D* = |*c*_2,0_| */c*_0,0_. **(B)** Histograms of deformation *D* at three developmental stages, revealing the dynamic nature of the process. **(C)** Top view of the three developmental stages with color-coded deformation. The color code relates to the histograms in **(B)**, where red nuclei are highly deformed. **(D)** Radial distributions of deformations *D* (upper), its binned statistics (middle) and nuclei count (bottom) over the three developmental stages.

Figure 4B shows how the percentage of highly deformed nuclei changes over the course of development. In general, it seems like the distribution of deformation is slowly shifting towards higher deformations as a whole. The maximum percentage of nuclei with *D* > 0.2 appears however at 6.5 hpf, around the stage of shield formation in Zebrafish [36]. A visual inspection of the labelled pictures obtained from BeadBuddy in 4C does not seem to exhibit any particular distribution of the deformed nuclei. To validate this quantitatively, an analysis of the correlation between radial distance and deformation is performed, clearly showing a homogeneous distribution of deformation around the embryo for each timepoint. The evolution of the distributions also indicates how the average value of deformation increases over time. Furthermore, the accumulation of the highly deformed nuclei at the edge of the embryo (Figure 4D, 8 hpf) is consistent with the known thinning of the cellular layer surrounding the embryo yolk as epiboly progresses.

This type of analysis could be done for any kind of tissue and geometry, given the flexibility of BeadBuddy and its convenient batch analysis mode. The output data - such as segmentations, labelled pictures, and libraries of SH coefficients - require minimal post-processing for such visual or quantitative studies. This opens the possibility for fast dynamical analysis over space and time.

### 3.2 Stress measurement in a developing embryo

Taking the analysis from simple nuclear shape analysis towards stress inference, one can take advantage of the pre-computed stress solutions based on SHs included in BeadBuddy. For objects that are spherical in the absence of a force and whose material properties are known, the surface stresses are can be directly evaluated. For an exemplary analysis, polyacrylamide (PAA) beads were injected in a developing Zebrafish embryo and analyzed using the same principles as described above.

However, in this case one is not limited to the analysis of relative deformations, but can feed the SH coefficients into the stress solver for each bead. The starting point for such an analysis is shown in Figure 5.

**Fig. 5.**
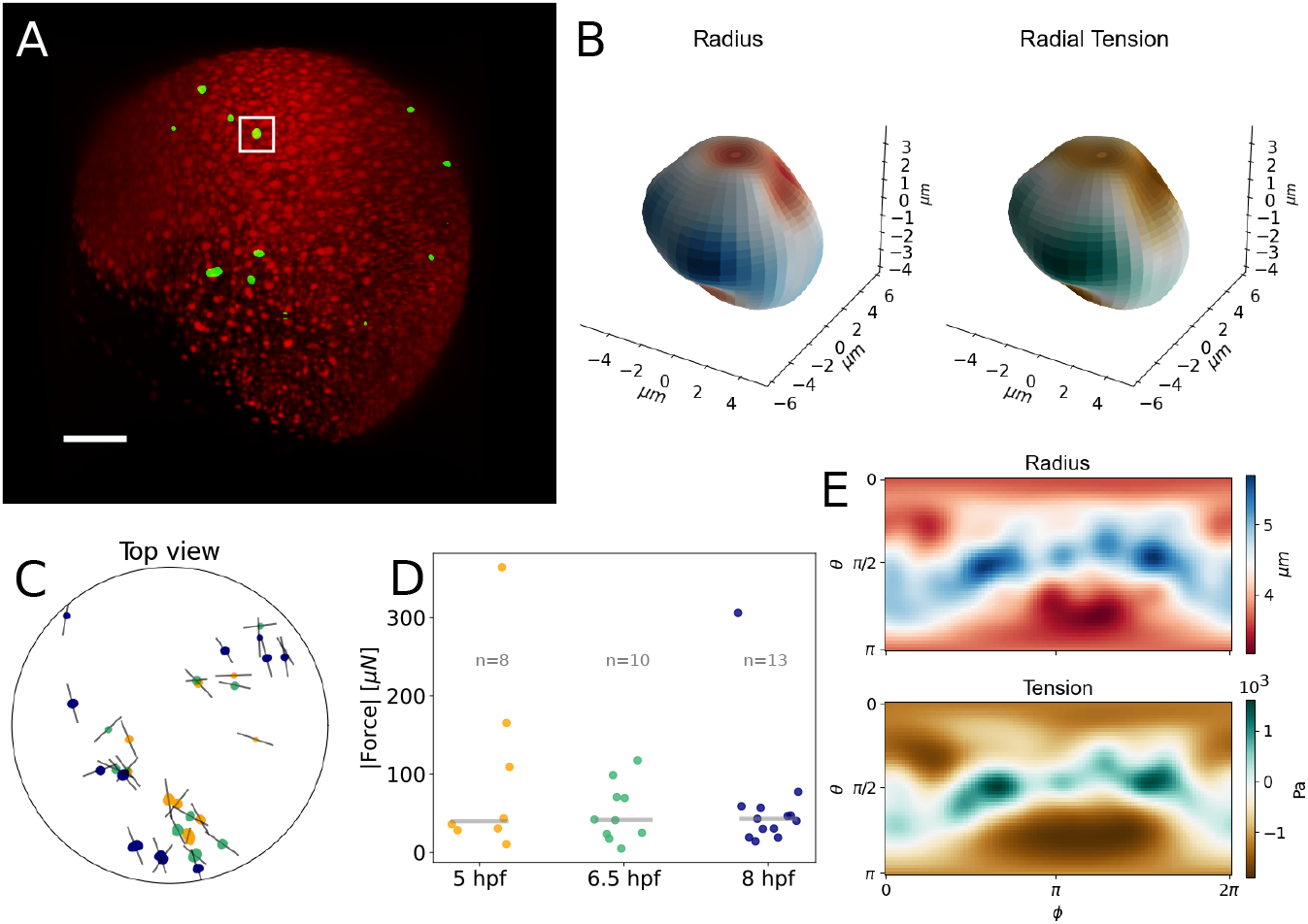
Batch analysis of PAA beads injected in a Zebrafish embryo. **(A)** Lightsheet image of the Zebrafish at 8 hpf, with H2AmCherry nuclei marked in red and PAA beads marked in green. Scalebar 100 µm. **(B)** 3D representations of an exemplary bead with color-coded radius (left) and radial tension (right). **(C)** Top view (animal pole) of the embryo with the population of measured PAA beads in three different developmental stages. The faint gray line crossing each bead represents their direction of maximum compression. **(D)** Measured forces acting on the surface of each bead in every timepoint. The grey horizontal line over each population represent its median. **(E)** Flattened *θ*-*ϕ* projections of the radius and radial tension shown in (B).

The stress solver renders deformation and radial stress maps in latitude-longitude (*θ-ϕ*) fashion (Figure 5E), as well as 3D projections over the deformed bead (Figure 5B). This kind of representation enables the latter exploration of correlations between the beads deformations and their surface interactions [29]. By numerically integrating the radial stress over the surface, the total force sensed by each bead can be calculated as well. An example of such a collective analysis is shown in Figure 5C,D. As shown in the previous section, the geometry of Zebrafish development facilitates the exploration of 3D data using different spherical projections. A top view, looking at the animal pole of the embryo, in Figure 5C shows the same collection of detected beads in three different stages of development. Their movement reveals a collective motion together with the blastoderm during epiboly, with beads in later stages being closer to the embryo edge. Thanks to the intrinsic deformation analysis via SHs, the main compression direction (grey line in Figure 5C) can be studied together with their position, which in this case does not seem to reveal any particular pattern. Thanks to the batch analysis mode in BeadBuddy, the total force felt by each bead at every timepoint can be calculated within seconds, enabling dynamics analysis as in Figure 5D. The median value of force sensed by a PAA bead in all timepoints was40 µN, distributed quite uniformly around the embryo.

### 3.3 Forces in reconstituted skeletal muscle

Any biophysical study at the cell or tissue level might benefit from the injection and analysis of force sensors. In some particular cases however, the mechanical information has proven to be key in the understanding of particular biological functions [37–39]. Such is the case of the homeostatic state of reconstituted skeletal muscle in healthy and diseased conditions [13, 40]. The usage of BeadBuddy has the potential to expand the analysis possibilities of such studies, revealing the main mechanical traits of a tissue. For this last example analysis in this article, we grew reconstituted skeletal muscle with PAA beads (7.8 kPa) embedded between two flexible posts, as shown in Figure 6A. After segmenting and calculating the forces sensed by each bead (*l*_max_ = 5), a series of conclusions can be drawn. The force value reveals a stronger force in beads which were in the vicinity of posts (where the muscle might be stretched the most), and less force in the central part of the tissue, as seen in Figure 6B (left). The difference between these two parts of the tissue is even clearer when looking at the distribution of the main compression direction, see Figure 6B (right).

**Fig. 6.**
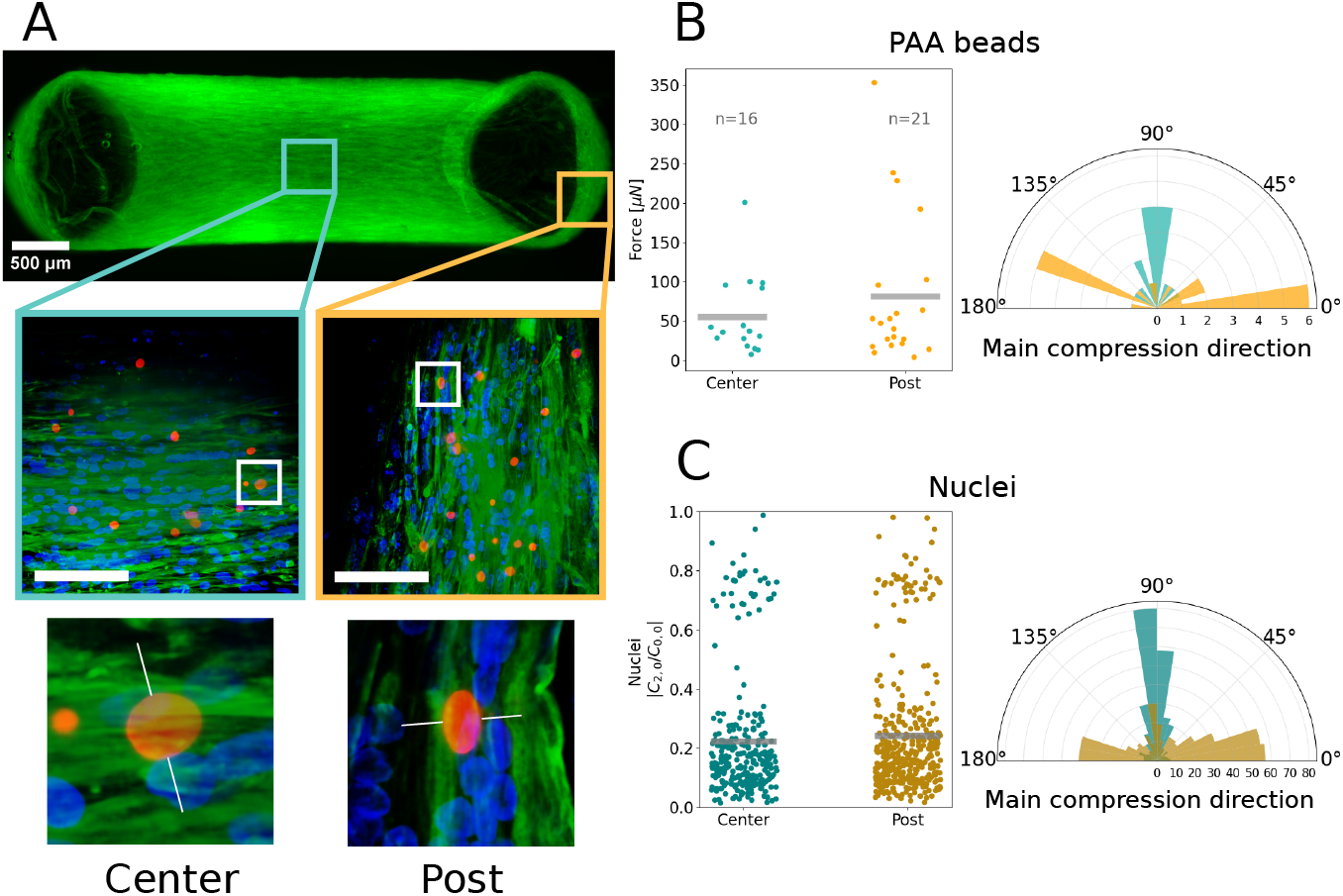
Analysis of forces and deformations inside a reconstituted skeletal muscle tissue via PAA beads and nuclei shape evaluation. **(A)** Confocal microscopy image of a human reconstituted skeletal muscle stained with Phalloidin in green, Hoechst in blue and PAA beads in red. A zoom window reveals details of the bead distribution (middle). Scalebar 100 µm. A further zoom focuses on a single bead in the Center and Post region, with a faint white line indicating the main compression direction of each bead. **(B)** Force measurements as felt by the PAA bead in the Center and Post region (left). Polar histogram of main compression directions in each region (right). **(C)** Similarly, distribution of deformation *D* = |*c*_2,0_| */c*_0,0_ of the nuclei in both regions, and histogram of compression direction.

In contrast, beads embedded in the Post regions show a main compression direction which is mostly horizontal, in opposition to the beads found in the Center region, exhibiting vertical compression. This clearly reveals how the beads feel the muscle fibers around them, being compressed perpendicularly to the fiber direction, and under a stronger load in the post regions, around which the fibers have to wrap to maintain the structural integrity of the muscle.

As in the previous case, the analysis is not limited to PAA beads but can be carried out likewise for all the nuclei present in the tissue. While nuclei have unknown mechanical parameters, they carry a higher statistical power due to their number. In this sense, a force cannot be associated to each nucleus, but one can resort to the calculation of the deformation parameter D instead (see Eq. (10). Nuclei in the Center region are less deformed than in the Post region, correlated to the distribution of forces in PAA beads. Furthermore, the distribution of the main compression direction perfectly mimics that of the beads. Nuclei seem to be mainly deformed perpendicularly to the direction of the muscle fibers, as if constrained by the geometry of the tissue.

## 4 Discussion

Here we have introduced a ready-to-use tool for the analysis of *in-vitro* or *in-vivo* elastic force sensors, which eliminates a great barrier between the experiments and straightforward, fast results. The cumbersome and computational-heavy methods found until now in the literature to tackle this task seem to have been a bottleneck for the analysis of data, either because of their complexity or their hindering slowness. By combining pre-computed analytical solutions to the elastic problem in the space of Spherical Harmonics and a user-friendly interface, we open the door for more biophysics labs to start exploiting the possibilities these shape and stress analyses offer. Elastic force sensors can be manufactured in a wet lab with relative easiness following already published protocols [13, 15, 18]. This, together with BeadBuddy, can be an addition to the toolbox of researchers dealing with the analysis of biophysical magnitudes at the cell and tissue level.

All of the necessary steps for the image analysis pipeline are embedded in the software, meaning that cropping, thresholding, segmenting and labelling of microscopy pictures can be achieved with several clicks, just before calculating the stress and deformation associated to each elastic sensor. The output of the analysis is also structured so that any post-process, when desired, can be carried out easily. We achieve this by providing the user with the SH coefficients, the main deformation axes, segmented and labelled pictures, and a collection of functions (`BeadBuddy TOOLS.py`) to plot and further analyze the data. Most of the figures presented in this articles have been generated using that toolbox, which demonstrates how quantitative data can be extracted from the experiments with minimal effort.

We have illustrated how this approach can be used to investigate the role of mechanics in different scenarios, either by exploring the time dependence of forces and deformation, or by studying spatial correlations which can give rise to a better understanding of the tissue structure. The analysis is not limited to elastic sensors, but could be extended, as shown, even to the study of cell nuclei. Even though the mechanical properties of nuclei can change during the cell cycle and according to the cell function, the study of nuclear deformation is a fundamental problem which can be simplified with BeadBuddy.We hope for our software to become a staple in the standard toolbox of any modern biophysics lab.

## 5 Materials and methods

### 5.1 Fabrication of polyacrylamide beads

PolyAcrylAmide (PAA) beads were prepared following a water-in-oil emulsion protocol. The water phase (AB mix) consisted of 500 µl of acrylamide solution (40 % v/v, Sigma, St. Louis, USA), 250 µl of N,N’-methylenebisacrylamide solution (2 % v/v, Sigma, St. Louis, USA) and 4 µl acrylic acid (99 % v/v, Sigma, St. Louis, USA). The tuning of the final mechanical properties of the beads is achieved by diluting this premix into 65% phosphate-buffered saline PBS (see recipe Table for exact ratios). The solution is finished by adding 6M NaOH to neutralize its pH. Right before the fabrication, the diluted AB mix is degassed in a vacuum chamber set at 50 mbar for a total of 10 minutes. After degassing, the free-radical cross-linking polymerization in the premix is initiated by adding a 10% ammonium persulphate solution (APS, AppliChem, Darmstadt, Germany).

**Table 1.**
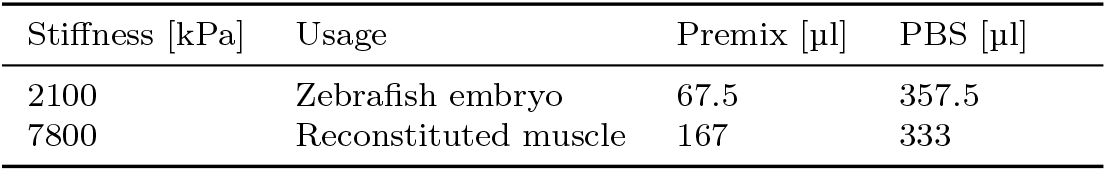
PAA beads mixture recipes.

Immediately after that, the water phase is transferred into an Eppendorf tube containing the oil phase of the emulsion, which consists of a solution of 3% Span80 (Sigma, St. Louis, USA) in n-Hexane (Merck, Darmstadt, Germany). A very fine emulsion is then achieved by sucking the dense water phase from the bottom of the tube and reinjecting it rapidly into the oil phase with a thin Hamilton syringe (Hamilton, Reno, USA). When the desired dispersion has been achieved, polymerization is catalyzed via addition of 3% (v/v) N,N,N’,N’-tetramethylethylenediamine (TEMED, Sigma, St. Louis, USA) and a further round of degassing at 50 mbar for additional 6 minutes. The supernatant is then discarded and the emulsion is to be kept at 85 ^°^C for 10 minutes to reach the proper PAA gelation. A day after, PAA beads are fully polimerized and can be dyed with different ATTO-NHS dyes (Atto-Tech, New York, USA) incubating them in solution for at least 30 minutes at room temperature.

### 5.2 Preparation of zebrafish embryos

Tg(βactin:H2AmCherry) transgenic embryos expressing mCherry fluorophore in all nuclei were raised at physiological conditions at 28 ^°^C in E3 medium. When they reached 256-cell stage (2.5 hpf), embryos were injected with a solution of PAA beads diluted in PBS. The injection was performed in the center of the blastoderm using gentle pressure through a custom pulled glass needle and an air pump injector (PV 820, World Precision Instruments, eject pressure 20 psi). At 4.5 hpf, the chorion surrounding the embryo was manually removed, and embryos were mounted in an imaging FEP tube loaded with 1% w/v low melting agarose. After some minutes, the agarose is stable enough to support the embryo in place, and soft enough to allow a normal embryonal development. Selective plane illumination microscopy was performed in a MuVi SPIM (Bruker) using 488 nm and 565 nm illumination wavelengths for PAA beads and the nuclei respectively. Four different orthogonal views with a step size of 2 µm were acquired and later fused to achieve an isotropic resolution of 1 µm^3^/voxel.

### 5.3 Reconstituted muscle tissue

Biomimetic human skeletal muscles were generated as previously described [13]. In brief, the human myoblast cell line AB1190, kindly provided by Vincent Mouly (Myoline platform of the Institut de Myologie, Paris, France) [41], was resuspended into an extracellular matrix mixture composed of DMEM (40 % v/v, Capricorn), 4 mg/ml bovine fibrinogen (Sigma) in 0.9 % (w/v) NaCl solution in water and Geltrex™ (20 % v/v, Gibco) at a concentration of 1.2 × 10^7^ cells per ml. Additionally, 1 × 10^4^ PAA beads (7.8 kPa, fluorescently labeled with Atto-565) per ml was added to the cell-ECM mixture. 0.5 units thrombin (Sigma) per mg of fibrinogen was used to initiate fibrin polymerization. Per tissue, 25 µl of cell-ECM mixture was casted into each well of the culture mold and subsequently incubated for 5 min at 37 ^°^C until the fibrin-based matrix polymerized. Skeletal Muscle Cell Growth medium (PROMOCELL) supplemented with 15 % fetal calf serum (FCS, Sigma), 1 % penicillin-streptomycin (Gibco) and 1.5 mg/ml 6-aminocaproic acid (ACA, Sigma) was then added to the 3D tissue culture and placed into a humidified incubator at 37 ^°^C and 5 % CO_2_. After two days, the media was changed to differentiation medium composed of DMEM supplemented with 2 % horse serum (HS, Sigma), 1 % penicillin-streptomycin (Gibco), 2 mg/ml ACA and 10 µg/ml human recombinant insulin (SAFC Biosciences). Differentiation media were changed every other day.

### 5.4 COMSOL calculations

Reference calculations using the finite element method were done using COMSOL Multiphysics. For these, the geometrically non-linear structural mechanics with linear elasticity was evaluated on imposed deformations in 3D, as given by the SH segmentation of recorded beads. Simple deformations were used to cross verify the correctness of both the analytical and this numerical approach. For concrete prediction matching of experimentally observed shapes, the mode representations were converted to explicit deformations on spheres with best-fit diameter. The radial component of the resulting surface traction could then be compared to predictions of the analytical calculations.

## 6 Code availability

The Python code of BeadBuddy is publicly available on https: https://gitlab.gwdg.de/betzlab_public/beadbuddy

## Appendix A BeadBuddy tutorial

BeadBuddy is contained in a Python script as found in our publicly available repository, and was developed for a Linux environment. After cloning the repository it is recommended to initialize an environment with the provided *env*.*ylm* file. Alternatively, the user can install all the necessary Python libraries manually before running the software for the first time in their machine.

The user interface can be called from a Linux terminal `python BeadBuddy.py` wherever a Python installation is available. After initialization, the user will be asked to select one of the available GPUs in their machine, and the software will run. The user can now load a 3D microscopy image in .*tiff* format to start the analysis. On the left-most display window the middle plane of the image will be shown, and the user can scroll through the z planes by moving the slider right below the image. Four segmentation parameters are offered to the user, namely:

**Fig. A1.**
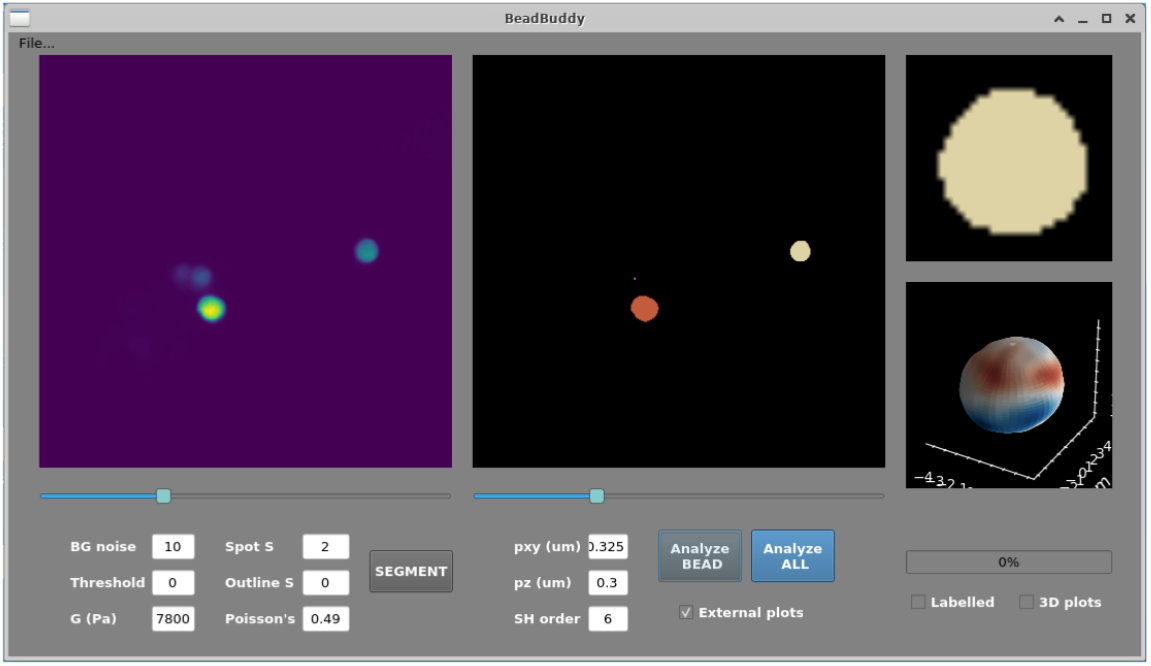
Screenshot of BeadBuddy in the process of analyzing the PAA beads embedded in muscle tissue, as shown in Figure 6.

- **BackGround noise**: the size (in pixel) of a Gaussian smoothing kernel to reduce noise in the image.
- **Threshold**: The pixel value from which the initial watershed algorithm will start detecting fluorescent signal.
- **Spot S**: The estimated size (in pixels) of the objects to be segmented.
- **Outline S**: The separation of the detected objects. The lower this value, the more fragmented the segmentation will be.

An initial set of parameters is suggested which has been proven useful as a starting point for good segmentations in a large variety of samples. By clicking on **SEGMENT**, the first segmentation will be performed on the fluorescence picture, and the results will be displayed in the middle window. Using its associated slider, the user can now explore the quality of the segmentation, for which the same plane on the original picture will be shown, as a direct comparison. The segmentation could be now refined to exclude big/small features, filter the possible noise associated with the image, split detections that lie in very close proximity, etc.

Once a satisfactory segmentation has been achieved, the user has the possibility of analyzing individual bodies or perform a batch analysis. To start with, the mechanical and acquisition parameters must be adjusted to the experimental needs: the shear modulus **G** and Poisson ratio ***ν*** of the material and the pixel height, width (**pxy**) and depth (**pz**) of the acquisition.

An individual body can be then analyzed by clicking on **Analyze BEAD** after clicking on the desired segmented body, and two new displays will be activated: a cross-section of the segmentation (top-right window) and a 3D reconstruction of the body with color-coded radius (bottom-right window). In case the checkbox **External plots** has been checked, two additional figures will be generated, as seen below. In case many bodies are present in the image and an analysis should be run on all of them, the batch mode can be started clicking on **Analyze ALL**. This will iterate through all the available bodies and save the following outputs in a folder created in the same path as the original image:

**Fig. A2.**
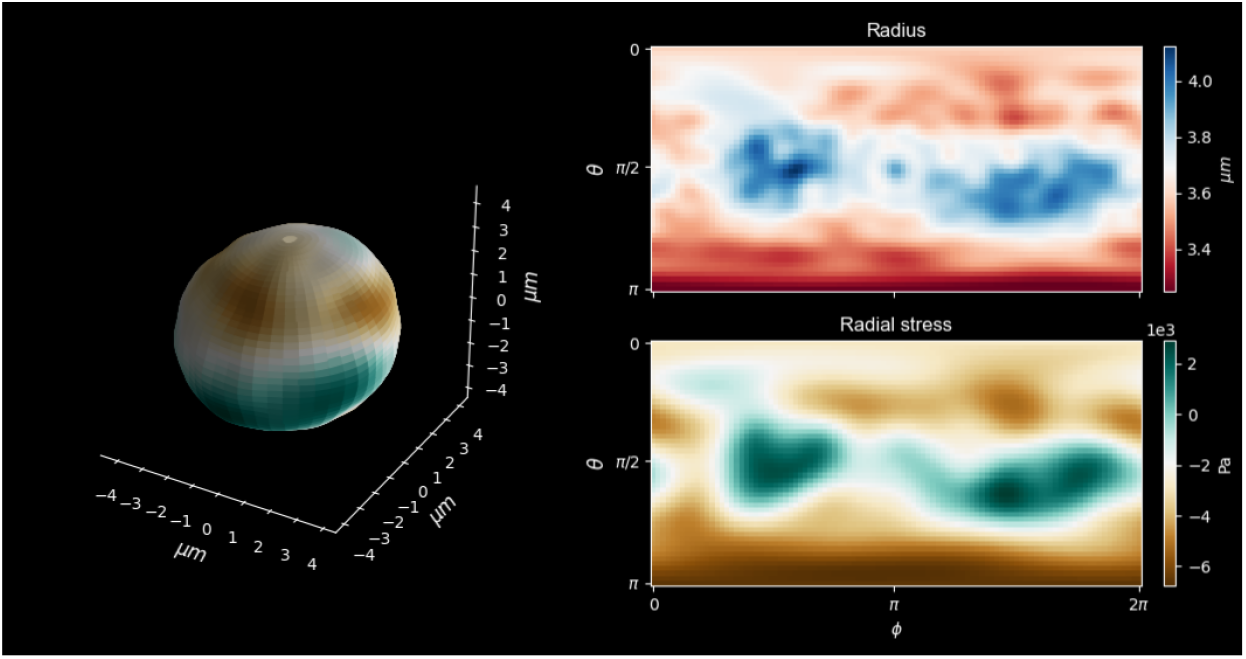
Additional figures obtained after clicking on the checkbox **External Plots**. On the left, color-coded radial tension as computed on the surface of the bead. On the right, plane projections of the radius (top) and radial tension (bottom), shown in *θ*-*ϕ* fashion.

- A Spherical Harmonics table associated to the expansion of the body surface.
- Two Euler angles [α_*x*_, α_*y*_] representing the deviation of the maximum compression direction of the body with respect to the vertical direction 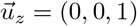.
- The segmented image, with the same dimensions as the original image.
- In case the corresponding checkboxes are ticked, the labelled image (with a single pixel value for each body) and the 3D reconstructions of each body (as .png figures) will also be stored.

Additionally a collection of post-processing functions is available in the repository which offers the user the possibility of batch-processing the output files and creating compelling and informative visualizations, such as the ones used in this article. The repository and the collection of post-processing scripts are still under expansion at the time of publication.

## Appendix B

Definition of Spherical Harmonics

Since many different conventions for Spherical Harmonics can be found in literature, we explicitly present here the one used for our analytical calculations, based on the work of Krüger et al. [25]. The normalized Spherical Harmonics are given by

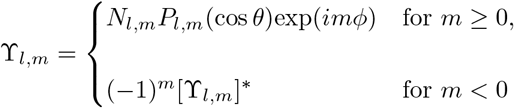

in the domain *θ* ∈ [− π/2, π/2], *ϕ* ∈ [™π/2, π/2]. The normalization factor *N*_*l,m*_ is given by

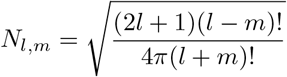

and *P*_*l,m*_ stands for the Legendre polynomials

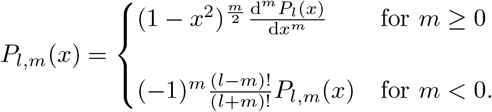

## Acknowledgements

This work was supported by the Deutsche Forschungsgemeinschaft Project ID: INST 186/1385-1 FUGG; 450595133. for T.B and A.J. R.W. is funded by the Deutsche Forschungsgemeinschaft (DFG, German Research Foundation) - WI 4170. The author would like to thank Kerstin von Roden for her invaluable help in the wet lab and during polymer fabrication, and the staff of the GZMB in Göttingen for their attention and care of the animals used in the study.

